## Supporting Tables for "A dual-reporter system for investigating and optimizing protein translation and folding in *E. coli*"

**Table S1. Primer sequences**

| Primer name | Primer sequence 5'- 3' |
|---|---|
| *GFP-fwd* | AGTATCTAGAATGCGTAAAGGAGAAGAACTT |
| *GFP-ASV-rev* | ACTGACTAGTTTAAACTGATGCAGCGTAGT |
| *GFP-mut3-rev* | ACTGACTAGTTTATTTGTATAGTTCATCCATGCC |
| *IbAp-fwd* | GAGCTTAATTAAAATTCATCTGTTGATCGTGG |
| *IbAp-rev* | GATATCTAGATAGCTCCTGAAATCAGCGAGAATGTAAG |
| *mCherry-fwd* | TCACCACCATCATTAGGATGGTGGTGATGATAATGGTTAGCAAAGGTGAAGAGGA |
| *mCherry-rev* | GGTGCTCGAGTGCGGCCGCAAGCTTTCATTTATACAGTTCATCCA |
| *BRCA1-BRCT-fwd* | ACTTTAAGAAGGAGATATACATATGGTCAACAAAAGAATGTCCATGGTG |
| *BRCA1-BRCT-rev2* | CACCATCCTAATGATGGTGGTGATGATGACTAGTCACAGGTGCCTCACACATCT |
| *PARP1-BRCT-fwd* | ACTTTAAGAAGGAGATATACATATGGTGAACTCCTCTGCT |
| *PARP1-BRCT-rev* | CACCATCCTAATGATGGTGGTGATGATGACTAGTTGGGGCCACAACTTCAACA |
| *E6-fwd* | ACTTTAAGAAGGAGATATACATATGGC |
| *P19-fwd* | ACTTTAAGAAGGAGATATACATATGCTG |
| *Coupling cassette-rev* | CACCATCCTAATGATGGTGGTGA |
| *NusA-fwd* | CGACATATGAACAAAGAAATTTTGGCTGTA GT |
| *NusA-rev* | ATACATATGGCTACCAGAGCCGCTACCCGCTTCGTCACCGAACCAG |
| *SUMO-fwd* | GACCATATGTCGGACTCAGAAGTCAAT |
| *SUMO-rev* | ATCCATATGGCTACCAGAGCCGCTACCACCACCAATCTGTTCTCTGT |
| *Mut-PARP1-BRCT-fwd* | GAGATATACATATGGTGAACTCCTCTGCT |
| *Mutagenesis-fwd* | CCCCTCTAGAAATAATTTTGTTTAACTTTAAGAAGGAGATATACATATG |
| *Mutagenesis-rev* | CCTAATGATGGTGGTGATGATGACTAGT |
| *Library-seq-fwd* | TCGTCGGCAGCGTCAGATGTGTATAAGAGACAGAACTTTAAGAAGGAGATATACATATG |
| *PARP1-BRCT-int-rev* | GTCTCGTGGGCTCGGAGATGTGTATAAGAGACAGTGAAGGCTCTTGGTGGAG |
| *PARP1-BRCT-int-fwd* | TCGTCGGCAGCGTCAGATGTGTATAAGAGACAGAACAAGGATGAAGTGAAGGC |
| *Library-seq-rev* | GTCTCGTGGGCTCGGAGATGTGTATAAGAGACAGATGATGGTGGTGATGATGACTA |
| *CI2-fwd* | ACTTTAAGAAGGAGATATACATATGAAGACAGAGTGGCCAGAGTTGGTGGGG |
| *CI2-rev* | CACCATCCTAATGATGGTGGTGATGATGGCCGACCCTGGGGACCTGGGCAATG |
| *CI2-V34G-fwd* | ATCATAGTTCTGCCGGGGGGGACAATTGTGACC |
| *CI2-V34G-rev* | GGTCACAATTGTCCCCCCCGGCAGAACTATGAT |
| *CI2-R48I-fwd* | ATATCGGATCGACCGCGTCATACTCTTTGTCGATAAACTC |
| *CI2-R48I-rev* | GAGTTTATCGACAAAGAGTATGACGCGGTCGATCCGATAT |
| *CI2-I57A-fwd* | CCTGGGGACCTGGGCAGCGTTGTCGAGTTTATCG |
| *CI2-I57A-rev* | CGATAAACTCGACAACGCTGCCCAGGTCCCCAGG |
| *CI2-F50A-fwd* | GACCGCGTCCGCCTCGCGGTCGATAAACTCGA |
| *CI2-F50A-rev* | TCGAGTTTATCGACCGCGAGGCGGACGCGGTC |
| *CI2-V63A-fwd* | CAGGTCCCCAGGGCCGGCCATCATCAC |
| *CI2-V63A-rev* | GTGATGATGGCCGGCCCTGGGGACCTG |

**Table S2. DNA and protein sequences for construction of vector and experimentally tested constructs**

| *Name* | *DNA sequence* | *Protein sequence* |
|---|---|---|
| *lbpAp* | AATTCATCTGTTGATCGTGGGTGTTGGCCTGATGAGTTATAGCGATCCCTTGCTGAAAATAACATCATCATTACGTCGCACTGTGGCGGCTATCGCACTTTAACGTTTCGTGCTGCCCCCTCAGTCTATGCAATAGACCATAAACTGCAAAAAAAAGTCCGCTGATAAGGCTTGAAAAGTTCATTTCCAGACCCATTTTTACATCGTAGCCGATGAGGACGCGCCTGATGGGTGTTCTGGCTACCTGACCTGTCCATTGTGGAAGGTCTTACATTCTCGCTGATTTCAGGAGCTA | |
| *GFP-ASV* | ATGCGTAAAGGAGAAGAACTTTTCACTGGAGTTGTCCCAATTCTTGTTGAATTAGATGGTGATGTTAATGGGCACAAATTTTCTGTCAGTGGAGAGGGTGAAGGTGATGCAACATACGGAAAACTTACCCTTAAATTTATTTGCACTACTGGAAAACTACCTGTTCCATGGCCAACACTTGTCACTACTTTCGGTTATGGTGTTCAATGCTTTGCGAGATACCCAGATCATATGAAACAGCATGACTTTTTCAAGAGTGCCATGCCCGAAGGTTATGTACAGGAAAGAACTATATTTTTCAAAGATGACGGGAACTACAAGACACGTGCTGAAGTCAAGTTTGAAGGTGATACCCTTGTTAATAGAATCGAGTTAAAAGGTATTGATTTTAAAGAAGATGGAAACATTCTTGGACACAAATTGGAATACAACTATAACTCACACAATGTATACATCATGGCAGACAAACAAAAGAATGGAATCAAAGTTAACTTCAAAATTAGACACAACATTGAAGATGGAAGCGTTCAACTAGCAGACCATTATCAACAAAATACTCCAATTGGCGATGGCCCTGTCCTTTTACCAGACAACCATTACCTGTCCACACAATCTGCCCTTTCGAAAGATCCCAACGAAAAGAGAGACCACATGGTCCTTCTTGAGTTTGTAACAGCTGCTGGGATTACACATGGCATGGATGAACTATACAAAAGGCCTGCAGCAAACGACGAAAACTACGCTGCATCAGTTTAA | MRKGEELFTGVVPILVELDGDVNGHKFSVSGEGEGDATYGKLTLKFICTTGKLPVPWPTLVTTFGYGVQCFARYPDHMKQHDFFKSAMPEGYVQERTIFFKDDGNYKTRAEVKFEGDTLVNRIELKGIDFKEDGNILGHKLEYNYNSHNVYIMADKQKNGIKVNFKIRHNIEDGSVQLADHYQQNTPIGDGPVLLPDNHYLSTQSALSKDPNEKRDHMVLLEFVTAAGITHGMDELYKRPAANDENYAASV |
| *GFP-mut3* | ATGCGTAAAGGAGAAGAACTTTTCACTGGAGTTGTCCCAATTCTTGTTGAATTAGATGGTGATGTTAATGGGCACAAATTTTCTGTCAGTGGAGAGGGTGAAGGTGATGCAACATACGGAAAACTTACCCTTAAATTTATTTGCACTACTGGAAAACTACCTGTTCCATGGCCAACACTTGTCACTACTTTCGGTTATGGTGTTCAATGCTTTGCGAGATACCCAGATCATATGAAACAGCATGACTTTTTCAAGAGTGCCATGCCCGAAGGTTATGTACAGGAAAGAACTATATTTTTCAAAGATGACGGGAACTACAAGACACGTGCTGAAGTCAAGTTTGAAGGTGATACCCTTGTTAATAGAATCGAGTTAAAAGGTATTGATTTTAAAGAAGATGGAAACATTCTTGGACACAAATTGGAATACAACTATAACTCACACAATGTATACATCATGGCAGACAAACAAAAGAATGGAATCAAAGTTAACTTCAAAATTAGACACAACATTGAAGATGGAAGCGTTCAACTAGCAGACCATTATCAACAAAATACTCCAATTGGCGATGGCCCTGTCCTTTTACCAGACAACCATTACCTGTCCACACAAT | MRKGEELFTGVVPILVELDGDVNGHKFSVSGEGEGDATYGKLTLKFICTTGKLPVPWPTLVTTFGYGVQCFARYPDHMKQHDFFKSAMPEGYVQERTIFFKDDGNYKTRAEVKFEGDTLVNRIELKGIDFKEDGNILGHKLEYNYNSHNVYIMADKQKNGIKVNFKIRHNIEDGSVQLADHYQQNTPIGDGPVLLPDNHYLSTQSALSKDPNEKRDHMVLLEFVTAAGITHGMDELYK |

| | | |
|---|---|---|
| | CTGCCCTTTCGAAAGATCCCAACGAAAAGAGAGACCAC<br>ATGGTCCTTCTTGAGTTTGTAACAGCTGCTGGGATTACA<br>CATGGCATGGATGAACTATACAAATAA | |
| *Translation coupling cassette* | ACTAGTCATCATCACCACCATCATTAGGATGGTGGTGAT<br>GATA | |
| *mCherry* | ATGGTTAGCAAAGGTGAAGAGGATAATATGGCCATCAT<br>CAAAGAATTTATGCGCTTTAAAGTGCACATGGAAGGTA<br>GCGTTAATGGCCATGAATTTGAAATTGAAGGTGAAGGC<br>GAAGGTCGTCCGTATGAAGGCACCCAGACCGCAAAACT<br>GAAAGTTACCAAAGGTGGTCCGCTGCCGTTTGCATGGG<br>ATATTCTGAGTCCGCAGTTTATGTATGGTAGCAAAGCCT<br>ATGTTAAACATCCGGCAGATATCCCGGATTATCTGAAA<br>CTGAGCTTTCCGGAAGGTTTTAAATGGGAACGTGTGAT<br>GAATTTTGAAGATGGTGGTGTGGTGACCGTTACCCAGG<br>ATAGCAGCCTGCAGGATGGTGAATTTATCTATAAAGTT<br>AAACTGCGTGGCACCAATTTTCCGAGTGATGGTCCGGT<br>TATGCAGAAAAAAACAATGGGTTGGGAAGCAAGCAGC<br>GAACGTATGTATCCGGAAGATGGCGCACTGAAAGGTG<br>AAATTAAACAGCGCCTGAAACTGAAAGATGGTGGCCAT<br>TATGATGCAGAAGTTAAAACCACCTATAAAGCCAAAAA<br>ACCGGTTCAGCTGCCTGGTGCATATAACGTTAACATTAA<br>ACTGGATATCACCAGCCACAACGAGGATTATACCATTG<br>TTGAACAGTATGAACGTGCAGAAGGTCGCCATAGTACC<br>GGTGGTATGGATGAACTGTATAAATGA | MVSKGEEDNMAIIKEFMRFKVHMEGSVNG<br>HEFEIEGEGEGRPYEGTQTAKLKVTKGGPLPF<br>AWDILSPQFMYGSKAYVKHPADIPDYLKLSFP<br>EGFKWERVMNFEDGGVVTVTQDSSLQDGEF<br>IYKVKLRGTNFPSDGPVMQKKTMGWEASSE<br>RMYPEDGALKGEIKQRLKLKDGGHYDAEVKT<br>TYKAKKPVQLPGAYNVNIKLDITSHNEDYTIVE<br>QYERAEGRHSTGGMDELYK |
| *PARP1-BRCT* | GTGAACTCCTCTGCTTCAGCAGATAAGCCATTATCCAAC<br>ATGAAGATCCTGACTCTCGGGAAGCTGTCCCGGAACAA<br>GGATGAAGTGAAGGCCATGATTGAGAAACTCGGGGGG<br>AAGTTGACGGGGACGGCCAACAAGGCTTCCCTGTGCAT<br>CAGCACCAAAAAGGAGGTGGAAAAGATGAATAAGAAG<br>ATGGAGGAAGTAAAGGAAGCCAACATCCGAGTTGTGT<br>CTGAGGACTTCCTCCAGGACGTCTCCGCCTCCACCAAG<br>AGCCTTCAGGAGTTGTTCTTAGCGCACATCTTGTCCCCT<br>TGGGGGGCAGAGGTGAAGGCAGAGCCTGTTGAAGTTG<br>TGGCCCC | VNSSASADKPLSNMKILTLGKLSRNKDEVKA<br>MIEKLGGKLTGTANKASLCISTKKEVEKMNKK<br>MEEVKEANIRVVSEDFLQDVSASTKSLQELFL<br>AHILSPWGAEVKAEPVEVVAP |
| *BRCA1-BRCT-truncated* | GTCAACAAAAGAATGTCCATGGTGGTGTCTGGCCTGAC<br>CCCAGAAGAATTTATGCTCGTGTACAAGTTTGCCAGAA<br>AACACCACATCACTTTAACTAATCTAATTACTGAAGAGA<br>CTACTCATGTTGTTATGAAAACAGATGCTGAGTTTGTGT<br>GTGAACGGACACTGAAATATTTTCTAGGAATTGCGGGA<br>GGAAAATGGGTAGTTAGCTATTTCTGGGTGACCCAGTC<br>TATTAAAGAAAGAAAAATGCTGAATGAGCATGATTTTG<br>AAGTCAGAGGAGATGTGGTCAATGGAAGAAACCACCA<br>AGGTCCAAAGCGAGCAAGAGAATCCCAGGACAGAAAG<br>ATCTTCAGGGGGCTAGAAATCTGTTGCTATGGGCCCTT<br>CACCAACATGCCCACAGATCAACTGGAATGGATGGTAC | VNKRMSMVVSGLTPEEFMLVYKFARKHHITL<br>TNLITEETTHVVMKTDAEFVCERTLKYFLGIAG<br>GKWVVSYFWVTQSIKERKMLNEHDFEVRGD<br>VVNGRNHQGPKRARESQDRKIFRGLEICCYG<br>PFTNMPTDQLEWMVQLCGASVVKELSSFTL<br>GTGVHPIVVVQPDAWTEDNGFHAIGQMCE<br>APV |

| | | |
|---|---|---|
| | AGCTGTGTGGTGCTTCTGTGGTGAAGGAGCTTTCATCA<br>TTCACCCTTGGCACAGGTGTCCACCCAATTGTGGTTGTG<br>CAGCCAGATGCCTGGACAGAGGACAATGGCTTCCATGC<br>AATTGGGCAGATGTGTGAGGCACCTGTG | |
| *P19* | CTGCTGGAAGAAGTTCGCGCAGGCGATCGTCTGAGCG<br>GTGCAGCAGCACGTGGTGATGTTCAAGAAGTGCGTCGT<br>CTGCTGCATCGTGAACTGGTTCATCCTGATGCACTGAAT<br>CGTTTTGGTAAAACCGCACTGCAGGTTATGATGTTTGGT<br>AGCACCGCAATTGCACTGGAACTGCTGAAACAGGGTGC<br>AAGCCCGAATGTTCAGGATACCAGCGGCACCAGTCCGG<br>TTCATGATGCCGCACGTACCGGTTTTCTGGATACCCTGA<br>AAGTTCTGGTTGAACATGGTGCAGATGTTAATGTTCCG<br>GATGGTACAGGTGCACTGCCGATTCATCTGGCCGTGCA<br>AGAAGGTCATACCGCAGTTGTTAGCTTTCTGGCAGCAG<br>AAAGCGATCTGCATCGTCGTGATGCACGTGGTCTGACA<br>CCGCTGGAACTGGCACTGCAGCGTGGTGCACAGGATCT<br>GGTTGATATTCTGCAGGGTCACATGGTTGCACCGCTG | LLEEVRAGDRLSGAAARGDVQEVRRLLHREL<br>VHPDALNRFGKTALQVMMFGSTAIALELLKQ<br>GASPNVQDTSGTSPVHDAARTGFLDTLKVLV<br>EHGADVNVPDGTGALPIHLAVQEGHTAVVS<br>FLAAESDLHRRDARGLTPLELALQRGAQDLV<br>DILQGHMVAPL |
| *E6* | GCGCGCTTTGAGGATCCAACACGGCGACCCTACAAGCT<br>ACCTGATCTGTGCACGGAACTGAACACTTCACTGCAAG<br>ACATAGAAATAACCTGTGTATATTGCAAGACAGTATTG<br>GAACTTACAGAGGTATTTGAATTTGCATTTAAAGATTTA<br>TTTGTGGTGTATAGAGACAGTATACCGCATGCTGCATG<br>CCATAAATGTATAGATTTTTATTCTAGAATTAGAGAATT<br>AAGACATTATTCAGACTCTGTGTATGGAGACACATTGG<br>AAAAACTAACTAACACTGGGTTATACAATTTATTAATAA<br>GGTGCCTGCGGTGCCAGAAACCGTTGAATCCAGCAGA<br>AAAACTTAGACACCTTAATGAAAAACGACGATTCCACA<br>ACATAGCTGGGCACTATAGAGGCCAGTGCCATTCGTGC<br>TGCAACCGAGCACGACAGGAAAGACTCCAACGACGCA<br>GAGAAACACAAGTA | ARFEDPTRRPYKLPDLCTELNTSLQDIEITCVY<br>CKTVLELTEVFEFAFKDLFVVYRDSIPHAACHK<br>CIDFYSRIRELRHYSDSVYGDTLEKLTNTGLYN<br>LLIRCLRCQKPLNPAEKLRHLNEKRRFHNIAG<br>HYRGQCHSCCNRARQERLQRRRETQV |
| *NusA* | AACAAAGAAATTTTGGCTGTAGTTGAAGCCGTATCCAA<br>TGAAAAGGCGCTACCTCGCGAGAAGATTTTCGAAGCAT<br>TGGAAAGCGCGCTGGCGACAGCAACAAAGAAAAAATA<br>TGAACAAGAGATCGACGTCCGCGTACAGATCGATCGCA<br>AAAGCGGTGATTTTGACACTTTCCGTCGCTGGTTAGTTG<br>TTGATGAAGTCACCCAGCCGACCAAGGAAATCACCCTT<br>GAAGCCGCACGTTATGAAGATGAAAGCCTGAACCTGG<br>GCGATTACGTTGAAGATCAGATTGAGTCTGTTACCTTTG<br>ACCGTATCACTACCCAGACGGCAAAACAGGTTATCGTG<br>CAGAAAGTGCGTGAAGCCGAACGTGCGATGGTGGTTG<br>ATCAGTTCCGTGAACACGAAGGTGAAATCATCACCGGC<br>GTGGTGAAAAAAGTAAACCGCGACAACATCTCTCTGGA<br>TCTGGGCAACAACGCTGAAGCCGTGATCCTGCGCGAAG<br>ATATGCTGCCGCGTGAAAACTTCCGCCCTGGCGACCGC<br>GTTCGTGGCGTGCTCTATTCCGTTCGCCCGGAAGCGCG<br>TGGCGCGCAACTGTTCGTCACTCGTTCCAAGCCGGAAA<br>TGCTGATCGAACTGTTCCGTATTGAAGTGCCAGAAATC | NKEILAVVEAVSNEKALPREKIFEALESALATA<br>TKKKYEQEIDVRVQIDRKSGDFDTFRRWLVV<br>DEVTQPTKEITLEAARYEDESLNLGDYVEDQIE<br>SVTFDRITTQTAKQVIVQKVREAERAMVVDQ<br>FREHEGEIITGVVKKVNRDNISLDLGNNAEAV<br>ILREDMLPRENFRPGDRVRGVLYSVRPEARG<br>AQLFVTRSKPEMLIELFRIEVPEIGEEVIEIKAA<br>ARDPGSRAKIAVKTNDKRIDPVGACVGMRG<br>ARVQAVSTELGGERIDIVLWDDNPAQFVINA<br>MAPADVASIVVDEDKHTMDIAVEAGNLAQA<br>IGRNGQNVRLASQLSGWELNVMTVDDLQA<br>KHQAEAHAAIDTFTKYLDIDEDFATVLVEEGF<br>STLEELAYVPMKELLEIEGLDEPTVEALRERAK<br>NALATIAQAQEESLGDNKPADDLLNLEGVDR<br>DLAFKLAARGVCTLEDLAEQGIDDLADIEGLT<br>DEKAGALIMAARNICWFGDEAGSGSGS |

| | | |
|---|---|---|
| | GGCGAAGAAGTGATTGAAATTAAAGCAGCGGCTCGCGATCCGGGTTCTCGTGCGAAAATCGCGGTGAAAACCAACGATAAACGTATCGATCCGGTAGGTGCTTGCGTAGGTATGCGTGGCGCGCGTGTTCAGGCGGTGTCTACTGAACTGGGTGGCGAGCGTATCGATATCGTCCTGTGGGATGATAACCCGGCGCAGTTCGTGATTAACGCAATGGCACCGGCAGACGTTGCTTCTATCGTGGTGGATGAAGATAAACACACCATGGATATCGCCGTTGAAGCCGGTAACCTGGCGCAGGCGATTGGCCGTAACGGTCAGAACGTGCGTCTGGCTTCGCAGCTGAGCGGTTGGGAACTCAACGTGATGACCGTTGACGACCTGCAGGCTAAGCATCAGGCGGAAGCGCACGCAGCGATCGACACCTTCACCAAATATCTCGACATCGACGAAGACTTCGCGACTGTTCTGGTAGAAGAAGGCTTCTCGACGCTGGAAGAATTGGCCTATGTGCCGATGAAAGAGCTGTTGGAAATCGAAGGCCTTGATGAGCCGACCGTTGAAGCACTGCGCGAGCGTGCTAAAAATGCACTGGCCACCATTGCACAGGCCCAGGAAGAAAGCCTCGGTGATAACAAACCGGCTGACGATCTGCTGAACCTTGAAGGGGTAGATCGTGATTTGGCATTCAAACTGGCCGCCCGTGGCGTTTGTACGCTGGAAGATCTCGCCGAACAGGGCATTGATGATCTGGCTGATATCGAAGGGTTGACCGACGAAAAAGCCGGAGCACTGATTATGGCTGCCCGTAATATTTGCTGGTTCGGTGACGAAGCGGGTAGCGGCTCTGGTAGC | |
| *SUMO* | TCGGACTCAGAAGTCAATCAAGAAGCTAAGCCAGAGGTCAAGCCAGAAGTCAAGCCTGAGACTCACATCAATTTAAAGGTGTCCGATGGATCTTCAGAGATCTTCTTCAAGATCAAAAAGACCACTCCTTTAAGAAGGCTGATGGAAGCGTTCGCTAAAAGACAGGGTAAGGAAATGGACTCCTTAAGATTCTTGTACGACGGTATTAGAATTCAAGCTGATCAGACCCCTGAAGATTTGGACATGGAGGATAACGATATTATTGAGGCTCACAGAGAACAGATTGGTGGTGGTAGCGGCTCTGGTAGC | SDSEVNQEAKPEVKPEVKPETHINLKVSDGSSEIFFKIKKTTPLRRLMEAFAKRQGKEMDSLRFLYDGIRIQADQTPEDLDMEDNDIIEAHREQIGGGSGSGS |
| CI2 | ATGAAGACAGAGTGGCCAGAGTTGGTGGGGAAATCGGTGGAGGAGGCCAAGAAGGTGATTCTGCAGGACAAGCCAGAGGCGCAAATCATAGTTCTGCCGGTGGGGACAATTGTGACCATGGAATATCGGATCGACCGCGTCCGCCTCTTTGTCGATAAACTCGACAACATTGCCCAGGTCCCCAGGGTCGGC | MKTEWPELVGKSVEEAKKVILQDKPEAQIIVLPVGTIVTMEYRIDRVRLFVDKLDNIAQVPRVG |

**Table S3. Primers for construction of PARP1-BRCT mutants**

| Mutant | Fragment 1 | Fragment 2 |
|---|---|---|
| **G20V** | G20V-FWD (CTGACTCTCGTGAAGCTGTCCC)<br>+<br>pET22-mut-rev (CTCACGCTGTAGGTATCTCAGTTCG) | G20V- REV (GGGACAGCTTCACGAGAGTCAG)<br>+<br>pET22-mut-fwd (CGAACTGAGATACCTACAGCGTGAG) |
| **G20W** | G20W-FWD (ATCCTGACTCTCTGGAAGCTGTCCC)<br>pET22-mut-rev (CTCACGCTGTAGGTATCTCAGTTCG) | G20W- REV (GGGACAGCTTCCAGAGAGTCAGGAT)<br>+ |

| | | |
|---|---|---|
| | | pET22-mut-fwd (CGAACTGAGATACCTACAGCGTGAG) |
| **A31T** | A31T-FWD (GATGAAGTGAAGACCATGATTGAG)<br>+<br>pET22-mut-rev (CTCACGCTGTAGGTATCTCAGTTCG) | A31T-REV (CTCAATCATGGTCTTCACTTCATC)<br>+<br>pET22-mut-fwd (CGAACTGAGATACCTACAGCGTGAG) |
| **I33N** | I33N-FWD (GAAGGCCATGAATGAGAAACTCG)<br>+<br>pET22-mut-rev (CTCACGCTGTAGGTATCTCAGTTCG) | I33N-REV (CGAGTTTCTCATTCATGGCCTTC)<br>+<br>pET22-mut-fwd (CGAACTGAGATACCTACAGCGTGAG) |
| **I33T** | I33T-FWD (GAAGGCCATGACTGAGAAACTCG)<br>+<br>pET22-mut-rev (CTCACGCTGTAGGTATCTCAGTTCG) | I33T-REV (CGAGTTTCTCAGTCATGGCCTTC)<br>+<br>pET22-mut-fwd (CGAACTGAGATACCTACAGCGTGAG) |
| **G37E** | G37E-FWD (TGAGAAACTCGAGGGGAAGTTGA)<br>+<br>pET22-mut-rev (CTCACGCTGTAGGTATCTCAGTTCG) | G37E- REV (TCAACTTCCCCTCGAGTTTCTCA)<br>+<br>pET22-mut-fwd (CGAACTGAGATACCTACAGCGTGAG) |
| **G37R** | G37R-FWD (TGAGAAACTCCGGGGGAAGTTGA)<br>+<br>pET22-mut-rev (CTCACGCTGTAGGTATCTCAGTTCG) | G37R- REV (TCAACTTCCCCCGGAGTTTCTCA)<br>+<br>pET22-mut-fwd (CGAACTGAGATACCTACAGCGTGAG) |
| **G37W** | G37W-FWD (TGAGAAACTCTGGGGGAAGTTGA)<br>+<br>pET22-mut-rev (CTCACGCTGTAGGTATCTCAGTTCG) | G37W- REV (TCAACTTCCCCCAGAGTTTCTCA)<br>+<br>pET22-mut-fwd (CGAACTGAGATACCTACAGCGTGAG) |
| **G38R** | G38R-FWD (AGAAACTCGGGCGGAAGTTGAC)<br>+<br>pET22-mut-rev (CTCACGCTGTAGGTATCTCAGTTCG) | G38R-REV (GTCAACTTCCGCCCGAGTTTCT)<br>+<br>pET22-mut-fwd (CGAACTGAGATACCTACAGCGTGAG) |
| **C50Y** | C50Y-FWD (GCTTCCCTGTACATCAGCACCAA)<br>+<br>pET22-mut-rev (CTCACGCTGTAGGTATCTCAGTTCG) | C50Y-REV (TTGGTGCTGATGTACAGGGAAGC)<br>+<br>pET22-mut-fwd (CGAACTGAGATACCTACAGCGTGAG) |
| **S52I** | S52I-FWD (CCTGTGCATCATCACCAAAAAG)<br>+<br>pET22-mut-rev (CTCACGCTGTAGGTATCTCAGTTCG) | S52I- REV (CTTTTTGGTGATGATGCACAGG)<br>+<br>pET22-mut-fwd (CGAACTGAGATACCTACAGCGTGAG) |
| **S52N** | S52N-FWD (CTGTGCATCAACACCAAAAAGGAG)<br>+<br>pET22-mut-rev (CTCACGCTGTAGGTATCTCAGTTCG) | S52N- REV (CTCCTTTTTGGTGTTGATGCACAG)<br>+<br>pET22-mut-fwd (CGAACTGAGATACCTACAGCGTGAG) |
| **I72N** | I72N-FWD (GGAAGCCAACAACCGAGTTGTG)<br>+<br>pET22-mut-rev (CTCACGCTGTAGGTATCTCAGTTCG) | I72N- REV (CACAACTCGGTTGTTGGCTTCC)<br>+<br>pET22-mut-fwd (CGAACTGAGATACCTACAGCGTGAG) |
| **V74F** | V74F-FWD (CCAACATCCGATTTGTGTCTGAG)<br>+<br>pET22-mut-rev (CTCACGCTGTAGGTATCTCAGTTCG) | V74F- REV (CTCAGACACAAATCGGATGTTGG)<br>+<br>pET22-mut-fwd (CGAACTGAGATACCTACAGCGTGAG) |
| **V74I** | V74I-FWD (CCAACATCCGAATTGTGTCTGAG)<br>+<br>pET22-mut-rev (CTCACGCTGTAGGTATCTCAGTTCG) | V74I- REV (CTCAGACACAATTCGGATGTTGG)<br>+<br>pET22-mut-fwd (CGAACTGAGATACCTACAGCGTGAG) |
| **D78V** | D78V-FWD (TGTGTCTGAGGTCTTCCTCCAGG)<br>+<br>pET22-mut-rev (CTCACGCTGTAGGTATCTCAGTTCG) | D78V- REV (CCTGGAGGAAGACCTCAGACACA)<br>+<br>pET22-mut-fwd (CGAACTGAGATACCTACAGCGTGAG) |
| **Q81R** | Q81R -FWD (GACTTCCTCCGGGACGTCTCC)<br>+<br>pET22-mut-rev (CTCACGCTGTAGGTATCTCAGTTCG) | Q81R – REV (GGAGACGTCCCGGAGGAAGTC)<br>+<br>pET22-mut-fwd (CGAACTGAGATACCTACAGCGTGAG) |
| **A96P** | A96P-FWD (GTTGTTCTTACCGCACATCTTGTC)<br>+<br>pET22-mut-rev (CTCACGCTGTAGGTATCTCAGTTCG) | A96P- REV (GGACAAGATGTGCGGTAAGAACAAC)<br>+<br>pET22-mut-fwd (CGAACTGAGATACCTACAGCGTGAG) |
| **A96V** | A96V-FWD (GAGTTGTTCTTAGTGCACATCTTGTCC) | A96V- REV (GGACAAGATGTGCACTAAGAACAACTC) |

| | | |
|---|---|---|
| | +<br>pET22-mut-rev (CTCACGCTGTAGGTATCTCAGTTCG) | +<br>pET22-mut-fwd (CGAACTGAGATACCTACAGCGTGAG) |
| **H97L** | H97L-FWD (GTTCTTAGCGCTCATCTTGTCCC)<br>+<br>pET22-mut-rev (CTCACGCTGTAGGTATCTCAGTTCG) | H97L- REV (GGGACAAGATGAGCGCTAAGAAC)<br>+<br>pET22-mut-fwd (CGAACTGAGATACCTACAGCGTGAG) |
